# Osmolarity regulates *C. elegans* egg-laying behavior via parallel chemosensory and biophysical mechanisms

**DOI:** 10.1101/2024.12.30.630790

**Authors:** Emmanuel Medrano, Karen Jendrick, Julian McQuirter, Claire Moxham, Dominique Rajic, Lila Rosendorf, Liraz Stilman, Dontrel Wilright, Kevin M Collins

**Affiliations:** Department of Biology University of Miami 1301 Memorial Drive Coral Gables, FL 33146

**Keywords:** osmolarity, hydrostatic pressure, *C. elegans*, egg laying, circuit, behavior

## Abstract

Animals alter their behavior in response to changes in the environment. Upon encountering hyperosmotic conditions, the nematode worm *C. elegans* initiates avoidance and cessation of egg-laying behavior. While the sensory pathway for osmotic avoidance is well-understood, less is known about how egg laying is inhibited. We analyzed egg-laying behavior after acute and chronic shifts to and from hyperosmotic media. Animals on 400 mM sorbitol stop laying eggs immediately but then resume ∼3 hours later, after accumulating additional eggs in the uterus. Surprisingly, the hyperosmotic cessation of egg laying did not require known osmotic avoidance signaling pathways. Acute hyperosmotic shifts in hyperosmotic-resistant mutants overproducing glycerol also blocked egg laying, but these animals resumed egg laying more quickly than similarly treated wild-type animals. These results suggest that hyperosmotic conditions disrupt a ‘high-inside’ hydrostatic pressure gradient required for egg laying. Consistent with this hypothesis, animals adapted to hyperosmotic conditions laid more eggs after acute shifts back to normosmic conditions. Optogenetic stimulation of the HSN egg-laying command neurons in hyper-osmotic treated animals led to fewer and slower egg-laying events, an effect not seen following direct optogenetic stimulation of the postsynaptic vulval muscles. Hyperosmotic conditions also affected egg-laying circuit activity with the vulval muscles showing reduced Ca^2+^ transient amplitudes and frequency even after egg-laying resumes. Together, these results indicate that hyperosmotic conditions regulate egg-laying via two parallel mechanisms: a sensory pathway that acts to reduce HSN excitability and neurotransmitter release, and a biophysical mechanism where a hydrostatic pressure gradient reports egg accumulation in the uterus.

**Summary Statement:** We find that hyperosmotic conditions inhibit *C. elegans* egg laying through both a sensory pathway and a separate biophysical pathway affecting a ‘high-inside’ hydrostatic pressure gradient.

## Introduction

Like many other animals, *C. elegans* regulates behaviors like locomotion and egg laying in response to external environmental conditions (Sawin, 1996; Sawin et al., 2000; Yu et al., 2017; Zhu et al., 2014). Shifts to low external osmolarity promote *C. elegans* egg laying (Zhang et al., 2008), while high external osmolarity inhibits it (Sawin, 1996). How changes in external osmolarity mediate these effects remains unclear. Genetics screens for *C. elegans* mutants that affect osmotic avoidance have identified numerous genes required for the development and function of chemosensory neurons that mediate osmotic avoidance (Culotti and Russell, 1978). Nociceptive sensory neurons signal to inhibit egg laying in response to unfavorable environmental conditions, and thus, external osmolarity may similarly function through a sensory pathway to regulate egg-laying behavior. For example, elevated CO_2_ activates the BAG nociceptive neurons, driving release of neuropeptides which bind to the EGL-6 receptor expressed on the HSN command motor neurons which control egg laying (Fenk and de Bono, 2015; Ringstad and Horvitz, 2008). EGL-6 couples to Gα_o_ which signals through downstream effectors including IRK-1 to stabilize HSN electrical excitability (Emtage et al., 2012; Ravi et al., 2021), reducing serotonin release and vulval muscle contractility (Collins et al., 2016). However, it is unknown whether regulation of egg laying by external osmolarity is mediated through similar cellular and molecular mechanisms.

A study by Zhang et al. (2010) suggested the vulval uv1 neuroendocrine cells may mediate egg laying inhibition by elevated osmolarity. Like BAG, uv1 expresses neuropeptides which can inhibit egg laying, possibly through HSN-expressed receptors (Banerjee et al., 2017). Additionally, animals bearing a putative dominant-negative mutation for the OCR-2 TRPV channel subunit expressed in the uv1s (Jose et al., 2007) show hyperactive egg laying even under high osmolarity conditions. OCR-2 is expressed in additional sensory neurons where changes in OCR-2 channel function might alter egg laying in high osmolarity environments (Sokolchik et al., 2005; Tobin et al., 2002).

Hyperosmotic inhibition of egg laying may also be mediated by a sensory-independent pathway. Normally, under hyperosmotic stress, *C. elegans* shows a rapid (within minutes) decrease in relative body volume due to water loss to the surrounding environment. Genetics screens for mutant animals resistant to changes in osmolarity (Osr) have identified pathways that function internally to prevent the loss of water and hydrostatic pressure (Solomon et al., 2004; Wheeler and Thomas, 2006). For example, *osr-1* mutants display normal viability, size, and motility when grown in hyperosmotic environments (Solomon et al., 2004). Acclimatization to osmotic stress is accomplished by accumulation of glycerol that leads to an increase in relative body volume (Lamitina et al., 2004). *C. elegans* expresses various aquaporins that have been shown to increase water or glycerol permeability when exogenously expressed in *Xenopus* oocytes. Surprisingly, quadruple mutant worms of some of these aquaporins (*aqp-2, aqp-3, aqp-4,* and *aqp-8*) are able to survive hypotonic stress just as well as wild-type worms, suggesting other mechanisms of osmoregulation or compensatory mechanism (Huang et al., 2007). Worms maintain their internal pressure hydrostatically and are normally highly pressurized (∼140 kPa). Puncturing their cuticle with a fine needle leads to expulsion of internal organs (Gilpin et al., 2015; Avery and Thomas, 1997; Harris and Crofton, 1957). As such, osmotic regulation of egg laying may be mediated by the changes in internal pressure that accompany exposure to hyper- or hypoosmotic environments. These changes, in turn, regulate the ability of hydrostatic pressure to drive release of eggs into the environment following vulval muscle contraction.

We have previously shown that changes in internal stretch and pressure that accompany egg accumulation affect egg-laying circuit activity and behavior. Genetically or chemically sterilized animals lacking eggs have reduced Ca^2+^ activity in the HSN command neurons and the postsynaptic vulval muscles (Ravi et al., 2018b). Conversely, artificial induction of egg accumulation drives a homeostatic increase in HSN Ca^2+^ activity that drives release of the additional eggs (Ravi et al., 2018b). Feedback of egg accumulation can modulate the egg-laying vulval muscles independent of the HSNs (Collins et al., 2016; Ravi et al., 2021), suggesting the vulval muscles themselves are the ultimate target. Consistent with this, acute injection to mimic the stretch-dependent feedback of egg accumulation is sufficient to induce egg-laying circuit activity with the vulval muscles among the first to respond (Medrano and Collins, 2023). This suggests that changes in hydrostatic pressure resulting from changes in egg accumulation or osmolarity may facilitate changes in egg laying. Within a stretch model, hypoosmotic activation of egg laying has the same physical consequences as acute injection. Acute shifts of animals to hypoosmotic conditions drives a rapid influx of water into animals (Lamitina et al., 2004). The presumable increases in hydrostatic pressure inside the animal then promotes circuit activity and egg laying. Conversely, a decrease in internal hydrostatic pressure would be predicted to inhibit egg-laying circuit activity and behavior, as previously observed (Zhang et al., 2008; Zhang et al., 2010).

Here, we use manipulations of osmolarity to understand how sensory input and the hydrostatic pressure gradient control egg laying. To do this we place animals on solid media of high osmolarity and assay the consequences on egg laying. We find that egg laying is inhibited in both wild-type and numerous chemosensory and osmotic resistant mutant animals. Interestingly, we find that inhibition of egg laying by high osmolarity is acute, with animals resuming egg-laying behavior in hyperosmotic conditions after ∼3 hours. Shifting to lower osmotic conditions following chronic high osmolarity exposure drives an acute increase in egg laying, consistent with the hydrostatic pressure gradient stimulating egg release. We also find that optogenetic activation of the vulval muscles of animals in hyperosmotic conditions is sufficient to drive egg laying. Conversely, optogenetic stimulation of the HSNs in animals in the same conditions is insufficient, with animals showing reduced efficiency in opto-induced egg laying, a delay in egg-laying onset, as well as an increase in the duration of the egg-laying event. Furthermore, Ca^2+^ imaging of the vulval muscles shows an overall reduction in amplitude and frequency for animals exposed to high osmolarity conditions. Together, these results suggest hyperosmotic environment regulate egg-laying behavior through both biophysical and chemosensory pathways.

## Methods

### Nematode culture

*Caenorhabditis elegans* strains were maintained as hermaphrodites at 20 °C on Nematode Growth Medium (NGM) agar plates with *E. coli* OP50 as a source of food as described (Brenner, 1974). All strains are derived from the Bristol N2 wild-type strain. Behavior assays were performed using age-matched adult hermaphrodites ∼24-30 hours past the late L4 stage using recommended sample sizes (Chase and Koelle, 2004). A list of strains, mutants, and transgenes used in this study can be found in **Table 1**.

**Table 1:**
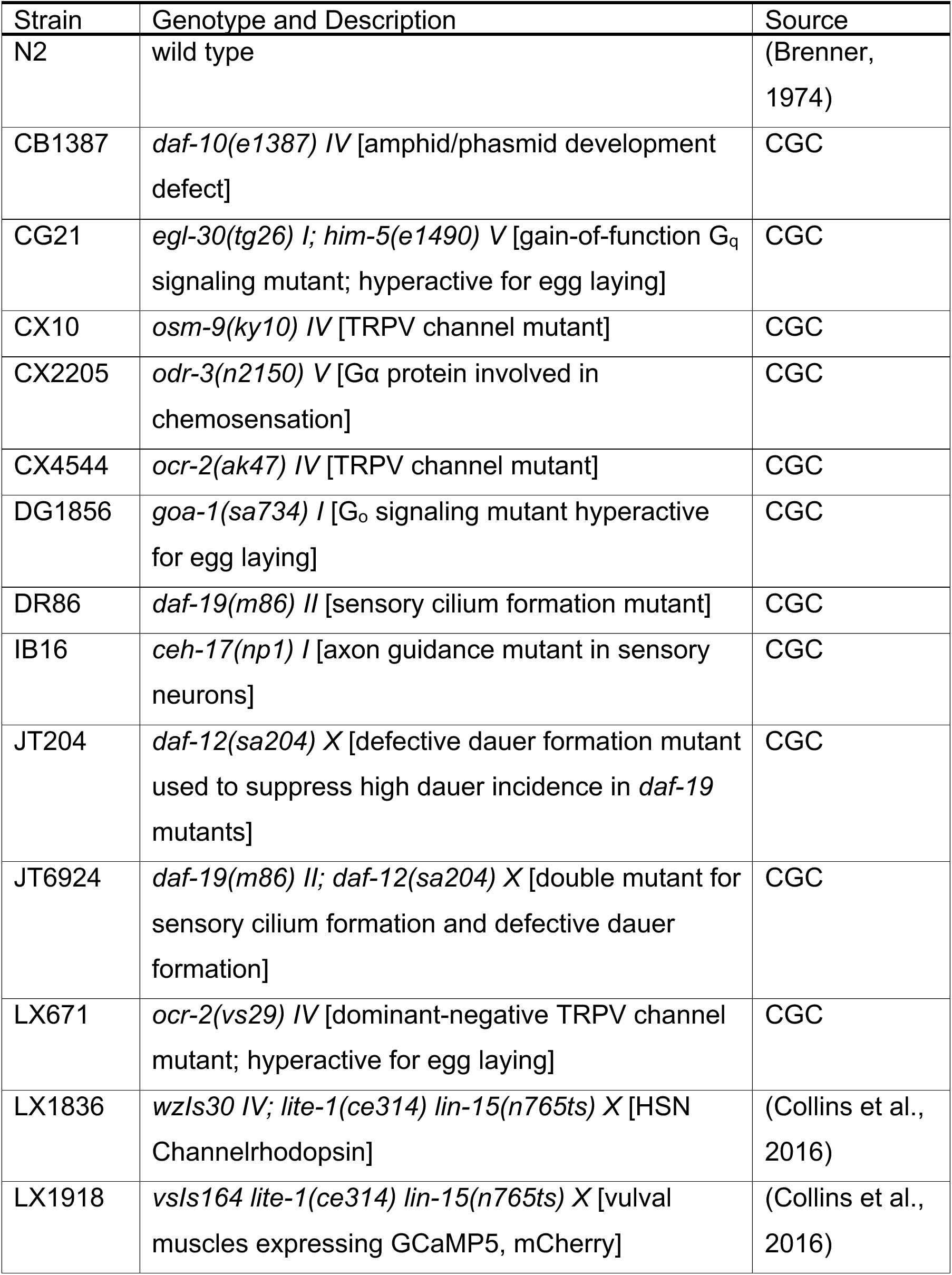
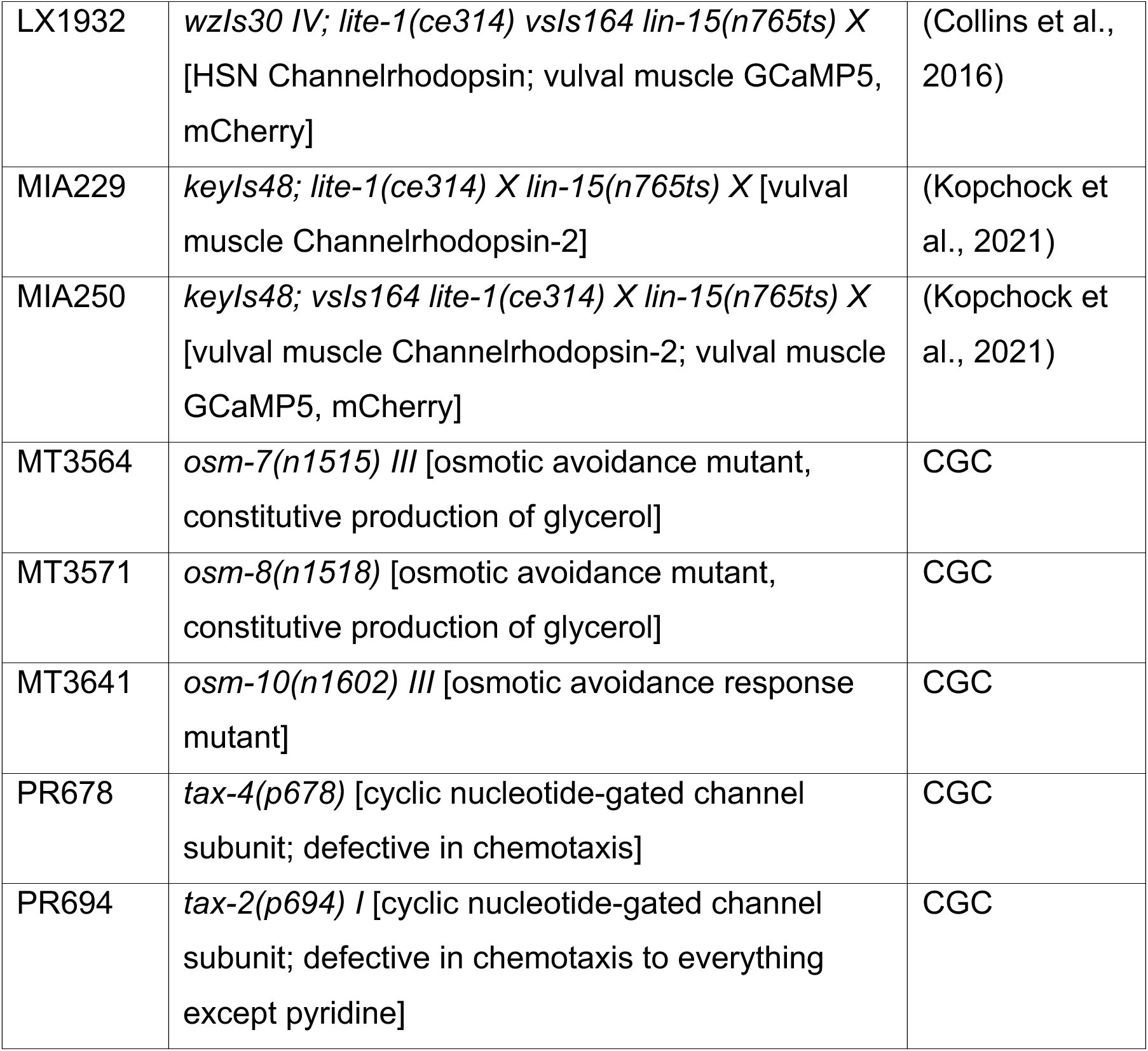
Strains used in this study.

### High osmolarity plates

Two different sets of high osmolarity plates were used in all experiments: 1.7% agar alone and NGM sorbitol plates. Agar plates were made as described (Alkema et al., 2005) adding acetic acid to a final concentration of 2 mM. Prior to solidifying, sorbitol was added from a 4M stock to either 100 mM or 400 mM final concentration. For long-term growth and shift experiments shown in **Figure 3**, NGM media contained 100 mM or 400 mM sorbitol added from a 4M stock solution. For the optogenetic experiments shown in **Figure 5**, NGM at ∼400 mOsm was prepared by adding sorbitol to 350 mM final concentration from a 4M stock. NGM alone (∼50 mOsm) was used as the ‘control’ media for these experiments. Agar or NGM media was then poured into 10-60 mm plates and allowed to dry for at least one day. All plates were seeded with OP50 bacteria. To allow for sufficient growth of bacteria, OP50 bacteria were concentrated to 10x through centrifugation. Plates were then seeded with 25 µL of 10x concentrated OP50 bacteria. Before all assays involving counting of eggs, 25 µL of 50 mM copper sulfate was added to the edge of the plate, swirled around the perimeter to form a ring of copper sulfate, discouraging animals from crawling up the sides of the plate where they would otherwise desiccate.

### Egg laying in high osmolarity

To assess how high osmolarity affects egg-laying behavior we placed 24-hr post L4 animals into agar plates containing either 100 mM or 400 mM sorbitol **(Figure 1A**). 3 – 5 animals were placed on replicates of each plate and allowed to lay eggs for 80 minutes. After this time, the total number of eggs laid by all animals was counted for each condition. The average number of eggs laid per animal was calculated by dividing the total number of eggs laid on a plate by the number of animals on that plate (typically 3-5). This same procedure was used in the candidate mutant screen to determine the effect of high osmolarity on the egg laying of these mutants. For chronic exposure experiments, laid eggs were counted every 30 minutes while animals were maintained on agar-sorbitol plates for up to 5.5 hours. Unlaid eggs were quantified as described (Chase and Koelle, 2004).

**Figure 1:**
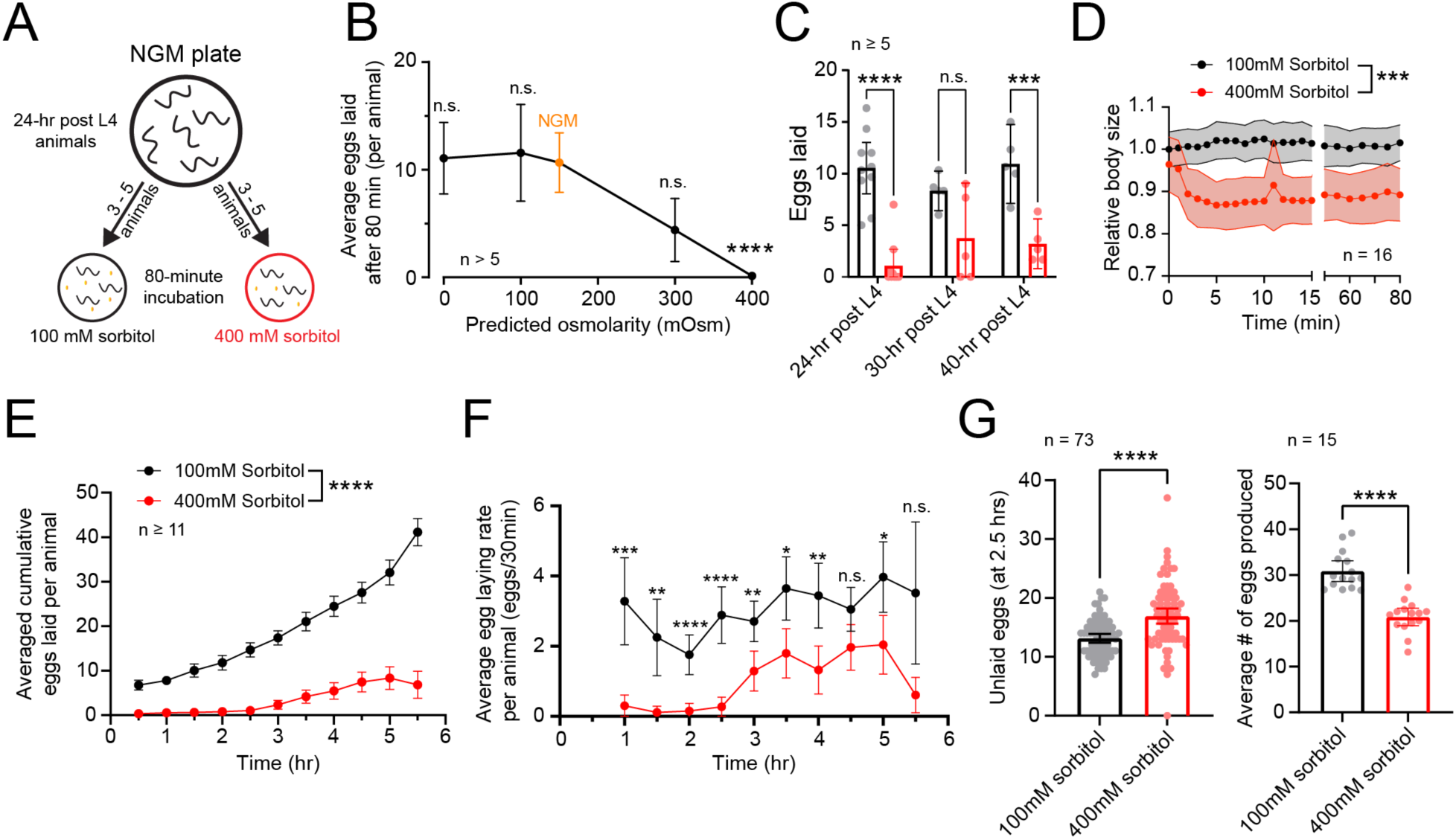
Hyperosmotic inhibition of egg laying behavior is restored after chronic exposure. **(A)** Diagram showing the assay for assessing egg laying in hyperosmotic conditions. Animals were placed into agar plates containing either 100 mM or 400 mM sorbitol. After an 80-minute period, eggs laid on the plate were counted. **(B)** Dose response curve showing average number of eggs laid after 80 minutes by animals on plates with different predicted osmolar concentrations. Orange dot represents average response of animals placed on NGM plates with an approximate osmolarity of 150 mOSm. Asterisks indicate p-value < 0.0001 (Kruskal-Wallis with Dunn’s multiple comparison test, all conditions compared to NGM as control). **(C)** Scatterplot with bar graphs showing mean number of eggs laid by 24-, 30-, or 40-hr post L4 animals on 100 mM (black) or 400 mM (red) sorbitol plates. Asterisks **** indicate p-value < 0.0001, *** indicate p-value < 0.001, n.s. indicates not significant (Two-way ANOVA with Bonferroni correction for multiple comparisons). **(D)** Relative body size over time of animals placed acutely onto 100 or 400 mM sorbitol plates. Asterisks *** indicate p-value < 0.001 (unpaired t-test) **(E)** Cumulative number of eggs laid over time by animals in sorbitol plates. Asterisks **** indicate p-value <0.0001 (Mixed-effects model with Bonferroni’s correction for multiple comparisons) **(F)** Number of eggs laid per each 30-minute interval from (E). Asterisks **** indicate p-value < 0.0001, *** indicates p-value < 0.001, ** indicates p-value < 0.01, * indicates p-value < 0.05, n.s. indicates not significant (Mixed-effects model with Bonferroni’s correction for multiple comparisons) **(G)** Bar graph with scatterplot showing number of unlaid eggs within the uterus of animals exposed to 100 mM or 400 mM sorbitol for a period of 2.5 hours (left). Average number of eggs produced by animals in either sorbitol conditions, obtained by combining the number of laid and unlaid eggs. All dots in data (B) – (G), except for (G left), represent the average response from multiple animals within a plate. Egg laying response from a plate was divided by the number of animals in the plate to obtain average response on a per animal basis. N equals the number of plates analyzed per condition, each bearing 3-5 worms. Error bars represent ± 95% confidence intervals for the mean.

### Relative body size estimate

To measure changes in relative body size after acute shifts to high osmolarity, eight animals were placed onto 100 mM or 400 mM agar plates. Animals were then recorded for 80 minutes using a FLIR Grasshopper 3 camera mounted on stereomicroscope (Leica M165 FC). Recordings were then analyzed on ImageJ to measure total body surface area. Measurements were performed every minute for the first 15 minutes and then every five minutes thereafter. All measurements were normalized by averaging the area of all eight animals during the first frame of the recording and subtracting this average from all other measurements to determine the normalized area.

### High osmolarity rescue experiments

To measure the effects of rescuing animals from high osmolarity on egg laying, animals were moved from 400 mM to 100 mM sorbitol plates. When animals were incubated for only 1 hour, sorbitol agar plates were used, but when animals were incubated on high osmolarity for one or more generations, the NGM sorbitol plates were used (described above). This was done to prevent any defects in development or changes in behavior that may accompany growth in non-nutrient growth media.

### Optogenetics

MIA229 and LX1932 strains expressing Channelrhodopsin-2 (ChR2) in either the vulval muscles or HSNs were used in optogenetic assays (see **Table 1**). Prior to optogenetic exposure, plates containing all-trans retinal (ATR) (Sigma Aldrich, R2500) were prepared by adding 200 µL of warmed OP50 bacteria in B Broth containing 0.4 mM ATR. Animals were typically maintained on ATR plates for multiple generations. A set of control animals were grown on plates lacking ATR. Animals were picked to NGM agar alone or NGM agar supplemented with 350 mM sorbitol (∼400 mOsm total) for 0.5- 1h prior to imaging. From these plates, an agar chunk was cut out and placed worms side down onto a 25×60mm glass coverslip for imaging, as previously described (Collins and Koelle, 2013; Ravi et al., 2018a). Worms were imaged through a 20x objective on a Zeiss Axio Observer.Z1 inverted compound microscope. Animals were exposed to a light off, on, off sequence that lasted 90 s. ChR2 was excited continuously at the 30 s time point for a 30 s duration by a 470 nm LED. An OTPG4 TTL Pulse Generator (Doric Instruments) was used to trigger a Grasshopper 3 camera (FLIR) which captured 2 x 2 binned, 1,024 x 1,024 8-bit JPEG brightfield image sequences of behavior at 75 frames per second. Egg laying was recorded during the whole 90 s recording. Measurements of time until first egg-event and duration of physical egg expulsion were performed in ImageJ from brightfield recordings by counting number of frames required for each event to occur. Duration of egg expulsion was measured from the moment the egg surface emerged just past the vulval opening until it was completely released into the environment.

### Vulval muscle Ca^2+^ imaging

Calcium imaging was performed in freely behaving LX1918, LX1932, and MIA250 animals (see Table 1) using a Zeiss Axio Observer microscope at 20x magnification, as previously described (Collins et al., 2016; Kopchock et al., 2021; Ravi et al., 2018a). Animals were exposed to 100 mM or 400 mM agar plates for a minimum of 60 minutes up to a maximum of 4.5 hours prior to Ca^2+^ imaging. Data was collected from animals alternating between treatment conditions (e.g. one animal from 100 mM sorbitol followed by the next from 400 mM sorbitol). Non-optogenetic LX1918 recordings (**Figure 4**) were typically 10 minutes long with time of osmotic exposure noted for recorded individuals. Optogenetic recordings (**Figure 6**) were typically 90 s with the same 470 nm blue light used to excite GCaMP5 fluorescence also serving to stimulate the co-expressed ChR2. A 10 msec on / 40 msec off duty cycle was used to trigger 470 and 590 LED illumination (Zeiss Colibri.2) and excite ChR2 and GCaMP5/mCherry fluorescence which was separated via a Hamamatsu Gemini W-View beam splitter and imaged by an ORCA Flash 4.0 V2 sCMOS camera at 20 frames per second. Sample recordings from animals grown without ATR were collected on the same day to ensure ATR-plus animals were showing a robust optogenetic response. Timing of egg-laying events were measured from synchronously recorded brightfield behavior data. Data from at least three independent experiments each containing multiple animals from all treatments were pooled for subsequent analysis.

### Statistical procedures

Results from independent experiments over several days were pooled (unless otherwise noted). Statistical analyses were performed using Prism version 8-10 (GraphPad). Isogenic control animals were always tested alongside mutants to account for possible batch effects. Student’s T-Test was performed in experiments with two conditions or genotypes when the data were normally distributed. In cases in which multiple genotypes or conditions were compared, the data were analyzed by one-way or two-way ANOVA. In cases in which data were found to be non-normal, a nonparametric equivalence test was performed (e.g., Mann-Whitney or Kruskal-Wallis). All tests were corrected for multiple comparisons (Bonferroni for ANOVA, Dunn for Kruskal-Wallis).

## Results

### *C. elegans* resumes egg-laying behavior after chronic exposure to hyperosmotic stress

To test osmotic inhibition of egg-laying we developed an assay in which animals were placed on solid media containing different concentrations of sorbitol. Sorbitol was chosen as a non-metabolic sugar that can induce the osmotic stress response in *C. elegans* (Chandler-Brown et al., 2015). After an 80-minute incubation period, we then counted the number of eggs laid (**Figure 1A**). We found a dose-dependent inhibition with media containing 400 mM sorbitol completely blocking egg laying. Treatment with 100 mM sorbitol, which approximates the osmolarity of Nematode Growth Media (NGM, ∼150 mOsm) (Yu et al., 2017), led to a similar number of eggs laid as animals on NGM (**Figure 1B**). The inhibition on elevated sorbitol media was not dependent on age, as older animals still showed a significant reduction in egg laying (**Figure 1C**). Consistent with prior studies (Lamitina et al., 2004), we also saw a ∼15% decrease in relative body size immediately following exposure to 400 mM sorbitol that persisted for at least one hour (**Figure 1D**).

While inhibition of egg laying via hyperosmotic exposure (Zhang et al., 2008) has been previously shown, we wondered whether *C. elegans* could acclimate and resume egg-laying after chronic exposure. To do this, we recorded egg laying every 30 minutes for a 5.5-hour period by animals placed on media containing either 100 or 400 mM sorbitol. While egg laying was initially blocked on 400 mM sorbitol, animals resumed egg laying after ∼2.5 hours **(Figure 1E**). This recovery was faster than the reported 24 hours required for acclimation to hyperosmotic environments via accumulation of glycerol (Lamitina et al., 2004). Although egg laying resumed, the rate of egg laying was lower from 400 mM- than 100 mM-treated animals during all 30-minute intervals (**Figure 1F**). Consequently, animals on 400 mM sorbitol media retained 17 eggs on average, which was significantly more than the 13 retained by animals on 100 mM sorbitol after 2.5 hours of exposure (**Figure 1G, left**). However, animals on 400 mM sorbitol also showed reduced egg production when accounting for the total number of eggs laid on plates and those retained within the uterus (**Figure 1G, right**). Together, these results show that hyperosmotic media temporarily inhibits egg laying in *C. elegans*. Egg laying resumes after 2.5 hours of hyperosmotic exposure, correlating with an increase in egg accumulation within the uterus. This resumption of egg laying supports our previous work in which increased stretch from egg accumulation serves as an additional signal to promote egg-laying active states (Medrano and Collins, 2023; Ravi et al., 2018b).

### Hyperosmolarity inhibits egg laying in chemosensory, osmotic avoidance, but not osmoregulatory mutants

To investigate the potential molecular and cellular mechanisms mediating osmolarity-driven regulation of egg-laying behavior, we screened a wide range of genetic mutants. Osmotic avoidance in *C. elegans* is mediated by the polymodal ASH neurons that mediate avoidance to mechanosensory and chemical stimuli (Bargmann, 2006; Hart et al., 1999; Kaplan and Horvitz, 1993). These neurons respond and signal through a wide range of molecules including heterotrimeric G proteins including ODR-3 (Roayaie et al., 1998), cyclic nucleotide-gated channels TAX-2/4, and TRPV channels OSM-9 and OCR-2 (Liedtke et al., 2003), among many others (Bargmann, 2006). Previous work has shown that 300 mOsm media fails to inhibit egg laying in *tax-2* or *tax-4* mutants with inhibition of egg laying in these mutants occurring only at higher (≥ 375 mOsm) osmolarities (Huang et al., 2023). Consistent with this, we found 400 mM sorbitol still inhibited egg laying in *tax-2*, *tax-4*, and other sensory defective mutants (**Figure 2A**). These results suggest mechanisms necessary for osmotic avoidance are not mediating the inhibition of egg laying by 400 mM sorbitol. To more completely inhibit chemosensation, we next tested mutants that affect cilia development, required generally for sensory neuron function (Bargmann, 2006; Swoboda et al., 2000) or axonal development (Pujol et al., 2000). As shown in **Figure 2B**, these mutants also showed inhibition of egg laying after our 80-minute exposure assay, further supporting the idea that sensory perception of high osmolarity is not required for osmotic inhibition of egg laying.

**Figure 2:**
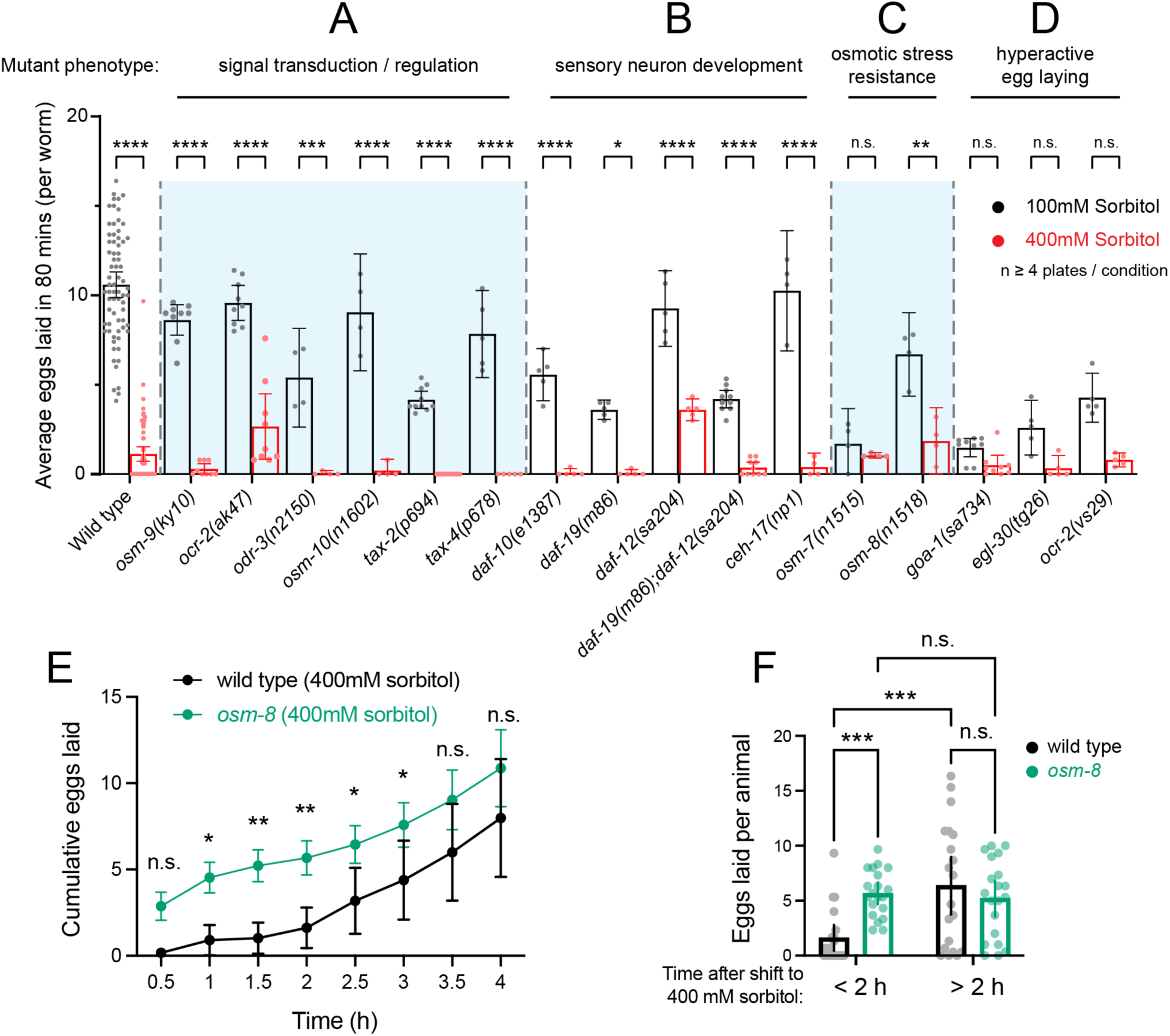
Mutants defective in osmosensation, sensory neuron development, the osmotic stress response, or hyperactive egg laying still show reduced egg laying in high osmolarity. **(A – D)** Scatterplot showing the average eggs laid within 80 minutes by wild type or mutants animals incubated in 100mM or 400mM sorbitol plates. Mutants are separated by category labels on the top and by the vertical dotted lines and shaded regions. Each dot represents the number of eggs laid by several animals in one plate divided by the total number of animals in that plate. (**E**) Time course of cumulative egg laying of wild-type (black) or *osm-8(n1518)* mutants (green) that over-produce glycerol after shifting to 400 mM sorbitol. Rate shown is the eggs laid per animal measured every 30 minutes averaged across 3 animals per plate from twenty independent biological replicates (plates) pooled from two experiments (ten each). (**F**) Average eggs laid by wild-type (black) or *osm-8* (green) mutant animals before (<2) and after (>2h) two hours of exposure to 400 mM sorbitol as shown in E. Asterisks **** indicate p-value <0.0001, *** indicate p-value <0.001, ** indicates p-value < 0.01, * indicates p-value <0.05, n.s. indicates not significant (Two-way ANOVA with Bonferroni correction for multiple comparisons). Error bars represent ± 95% confidence interval.

To determine if the osmotic stress response – in which animals accumulate glycerol to restore water loss – contributed to inhibition of egg laying, we next tested two mutants, *osm-7* and *osm-8*. These mutants show resistance to osmotic stress via constitutive production of glycerol, preventing the normal loss of hydrostatic pressure that accompanies hyperosmotic stress (Rohlfing et al., 2011; Wheeler and Thomas, 2006). *osm-7* mutants showed comparable levels of egg laying in 100 mM or 400 mM sorbitol, although we interpret this result as being caused by a low baseline level of egg laying in both conditions (**Figure 2C**). *osm-8* mutants had comparatively improved brood sizes but were still significantly inhibited for egg laying in the high osmolarity condition (**Figure 2C**). To test *osm-8* mutants further, we measured the time course of egg-laying inhibition and recovery after shifting to 400 mM sorbitol. *osm-8* mutants resumed egg laying within 1 hour, significantly earlier than wild-type control animals (**Figure 2E**), ultimately laying significantly more eggs within the first two hours (**Figure 2F**). These results suggest that elevation of internal glycerol to pre-emptively maintain a hydrostatic pressure gradient provides resistance to acute shifts to high sorbitol, allowing egg laying.

If high osmolarity inhibits activity in the egg-laying circuit, then mutants with increased Ca^2+^ activity may overcome this inhibition. To test this hypothesis, we tested hyperactive egg-laying mutants, specifically *goa-1(sa734*) mutants lacking inhibitory Gα_o_ signaling (Robatzek and Thomas, 2000) or *egl-30(tg26)* gain-of-function mutants with too much excitatory Gα_q_ signaling (Doi and Iwasaki, 2002). Both mutants have higher levels of egg-laying circuit activity and egg laying behavior on norm-osmolarity media (Dhakal et al., 2022; Ravi et al., 2021). However, egg laying in these mutants is still reduced on 400 mM sorbitol media (**Figure 2D**), although this is not significant, likely due to impaired egg production (Bastiani et al., 2003; Ségalat et al., 1995). *ocr-2(vs29)* mutant animals with altered TRPV channel function and hyperactive egg laying (Jose et al., 2007) were similarly inhibited by high osmolarity media (**Figure 2D**). Overall, these results suggest that sensory signaling is not strictly required for cessation of egg laying in response to hyperosmotic media exposure. This is supported by the wide range of mutants, defective in sensory neuron development or signaling, that still show inhibition by 400 mM sorbitol exposure. Our results from glycerol accumulation mutants suggest instead that physiological mechanisms that restore a hydrostatic pressure gradient allow egg-laying after shifts to high osmolarity. Together, these results support a biomechanical model in which a hydrostatic pressure gradient facilitates egg release.

### Recovery from hyperosmotic stress induces an acute increase in egg laying

We have previously shown that the egg-laying circuit induces a homeostatic increase in egg laying and in Ca^2+^ activity of the HSNs after artificial accumulation of unlaid eggs (Ravi et al., 2018b). Since hyperosmotic inhibition of egg laying leads to an increase in egg retention (**Figure 1G**), we next questioned whether animals incubated in high-osmolarity conditions would show an increase in egg-laying behavior once they were moved back to 100 mM sorbitol. As shown in **Figure 3A**, two control conditions were created: the 100 mM control from which animals were moved to a different set of 100 mM plates, and the 400 mM control condition from which animals were similarly moved from one set of 400 mM sorbitol plates to a different set. One experimental condition (400 mM → 100mM) involved moving animals incubated on 400 mM sorbitol for one hour to 100 mM plates (**Figure 3A**). Consistent with prior results (**Figure 1E**), 100 mM control animals showed a steady rate of egg laying despite the transfer while 100 mM animals transferred to 400 mM showed general inhibition of egg laying followed by resumption after 2.5 hours of exposure to hyperosmotic stress (**Figure 3B**). Animals that were switched from 400 to 100 mM sorbitol plates laid more eggs over time, ∼12 eggs on average per worm over 2 hours, compared to the ∼5-10 eggs from either group (**Figure 3B**). This result is consistent with a homeostatic rebound effect where an elevated ‘high inside’ hydrostatic pressure gradient is more favorable for egg laying immediately following the osmotic shift.

**Figure 3:**
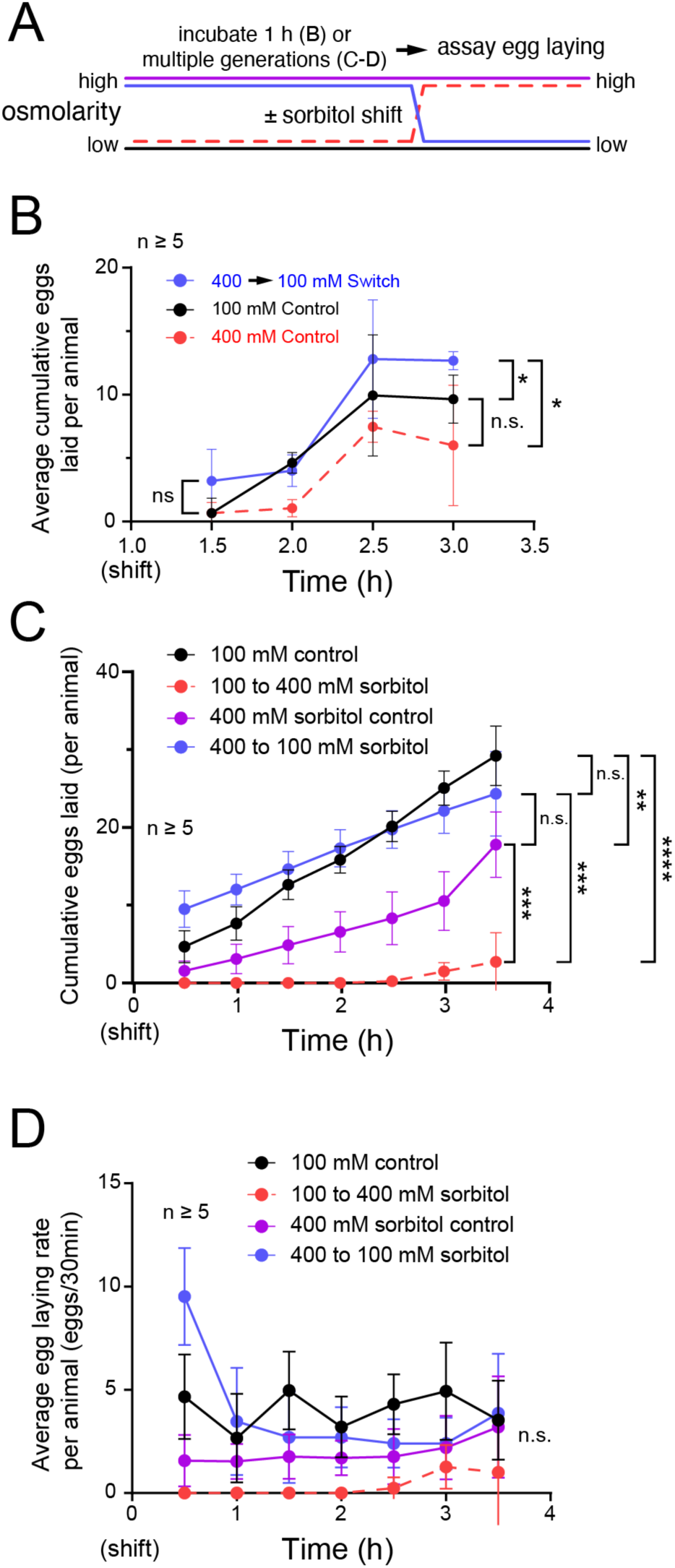
Rescue from hyperosmotic conditions induces a homeostatic increase in egg laying. **(A)** Diagram of experimental setup. Animals were incubated for either 1 hour on agar-sorbitol plates (B) or for multiple generations on NGM plates containing 100 mM or 400 mM sorbitol (C-D; see methods). Thereafter, animals were moved to either 100 mM or 400 mM agar-sorbitol plates (B) or NGM-sorbitol plates (C-D) and eggs laid on the new plates were counted every 30 minutes. **(B)** Cumulative number of eggs laid by animals incubated for 1 hour, or their entire lives **(C)**, followed a ‘shift’ to the indicated treatment plates (denoted by color). **(D)** Eggs laid per each 30-minute interval from (C). All dots in data (B) – (D) represent the average response from multiple animals within a plate. Egg laying response from a plate was divided by the number of animals in the plate to obtain average response per animal. Error bars represent ± 95% confidence interval. A mixed model statistical analysis with a Bonferroni correction for multiple comparisons was performed for each time series data. Due to the large number of comparisons, only significance at the beginning and end of each time series is shown. Only significance for the last timepoint is shown in (C) and (D). Asterisks **** indicate p-value <0.0001, *** indicate p-value <0.001, ** indicates p-value < 0.01, * indicate p-value <0.05, n.s. indicates not significant (p-value >0.05).

Since the maximal production of glycerol from acclimatization to osmotic stress takes ∼24 hours (Lamitina et al., 2004), we next tested if animals incubated on high sorbitol plates over multiple generations would show similar responses. Because these experiments require long-term incubations and growth that might not be optimal on simple agar plates, we grew worms on standard NGM supplement with either 100 mM or 400 mM sorbitol. If acclimatization to hyperosmotic environments is sufficient to prevent inhibition of egg laying, then we would expect no difference between animals grown on NGM media containing 100 or 400 sorbitol for their whole lives. We did find that animals shifted from NGM containing 400 mM sorbitol plates to the same media showed modestly lower levels of egg laying, reaching an average of ∼20 eggs laid per animal over 3.5 hours, compared to the ∼25 eggs laid per worm for the NGM plus 100 mM sorbitol control group (**Figure 3C**). This is significantly greater than the ∼3 eggs laid by animals shifted from NGM media with 100 mM sorbitol to NGM media plus 400 mM sorbitol. In contrast, animals grown on high sorbitol and down-shifted to 100 mM sorbitol showed a significant increase in egg laying for the first 30 minutes before declining to a similar steady-state level seen in 100 or 400 mM sorbitol treated control animals (**Figure 3D**). Overall, these results show that recovery from hyperosmotic stress induces an acute rebound in egg laying (< 30 min), consistent with a high-inside pressure gradient promoting egg release, followed by a return to normal rates of egg laying once the pressure gradient returns to normal. The low egg-laying rates maintained by animals kept on high osmolarity over multiple generations suggests a chronic inhibition of egg laying and/or germline activity that reduces egg laying even after the hydrostatic pressure gradient is restored.

### Vulval muscle Ca^2+^ activity does not recover following chronic hyperosmotic exposure

Acute exposure to high osmolarity has been shown to cause a decrease in HSN, VC, and vulval muscle Ca^2+^ peak frequency (Zhang et al., 2008; Zhang et al., 2010). If *C. elegans* can resume egg-laying behavior after chronic exposure to hyperosmotic environments, then presumably the egg-laying circuit shows resumption of activity. To test this, we recorded Ca^2+^ activity from the vulval muscles of animals incubated on 100 or 400 mM sorbitol plates for different amounts of time. Vulval muscle Ca^2+^ activity and egg-laying events were observed in both conditions (**Figure 4A**). However, animals on 100 mM plates laid eggs in 23% of recordings (8/35) while animals on 400 mM plates laid eggs in only 3% of all recordings (1/36), a significant difference (p-value =0.0333; Fisher’s exact test). Consistent with these lower rates of egg laying, hyperosmotic treatment yielded a decrease in average vulval muscle Ca^2+^ transient peak amplitude and frequency on a per animal basis (**Figure 4B and C**). Contrary to the observed resumption of egg-laying behavior (**Figure 1**), both vulval muscle Ca^2+^ transient amplitude and peak frequency did not show a positive correlation with time spent on high osmolarity (**Figure 4D and E**). Together, this data suggests that high osmolarity depresses vulval muscle Ca^2+^ activity while in a hyperosmotic environment, likely until they fully adapt. Accumulation of eggs may then serve to promote egg-laying behavior once animals reach a critical point of egg accumulation, as noted previously (Collins et al., 2016; Medrano and Collins, 2023).

**Figure 4:**
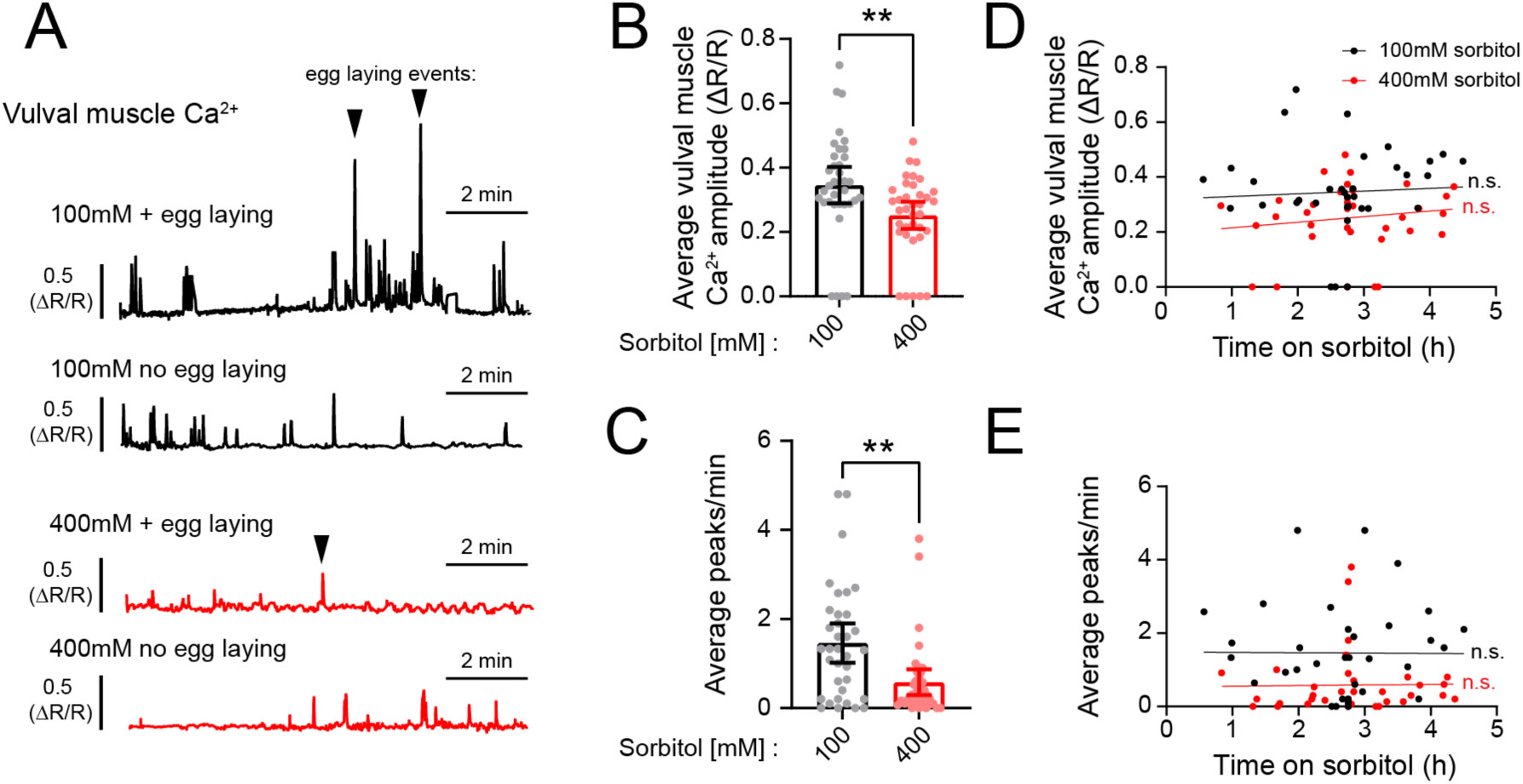
Vulval muscle Ca^2+^ activity remains inhibited after chronic exposure to high osmolarity. **(A)** Representative GCaMP5/mCherry fluorescence traces of animals exposed to NGM plates containing either 100mM (black) or 400mM (red) sorbitol. Traces show examples of animals in which no egg laying or at least one egg laying event was observed (denoted by triangle). **(B)** Average vulval muscle Ca^2+^ response from individual animals from both sorbitol conditions. Asterisks ** indicate p-value <0.01 (Mann-Whitney test). **(C)** Average Ca^2+^ frequency (peaks/minute) from both sets of animals. **(D – E)** Scatterplot with linear regression showing no correlation between average vulval muscle Ca^2+^ amplitude (D) or average Ca^2+^ frequency (E) and time exposed to sorbitol. n.s. represents no significance (simple linear regression).

### Hyperosmotic inhibition of egg laying is mediated in part by physical deficiencies in egg release

If hyperosmotic inhibition of egg laying is mediated by depression of circuit activity or a disruption of a pressure gradient required for proper expulsion of eggs, then animals forced to lay eggs in hyperosmotic conditions may show physical deficiencies depending on the way egg laying is induced. To test this hypothesis, we induced egg laying via optogenetic stimulation in transgenic animals expressing Channelrhodopsin-2 in the vulval muscles to bypass ongoing egg-laying circuit activity (Kopchock et al., 2021; Yan et al., 2024). Continuous blue-light exposure for 30 s led to egg laying in animals on control or high sorbitol plates (**Figure 5A**), with ∼80% of both groups showing light-induced egg laying (**Figure 5B**). We observed no deficiency in the amount of time taken for the first egg to be released (**Figure 5C**) or in the time required for eggs to be fully expelled from the uterus following optogenetic induction of vulval activity (**Figure 5D**). These results indicate that acute changes in the hydrostatic pressure gradient do not significantly alter egg release when driven in response to sustained optogenetic vulval muscle contraction.

**Figure 5:**
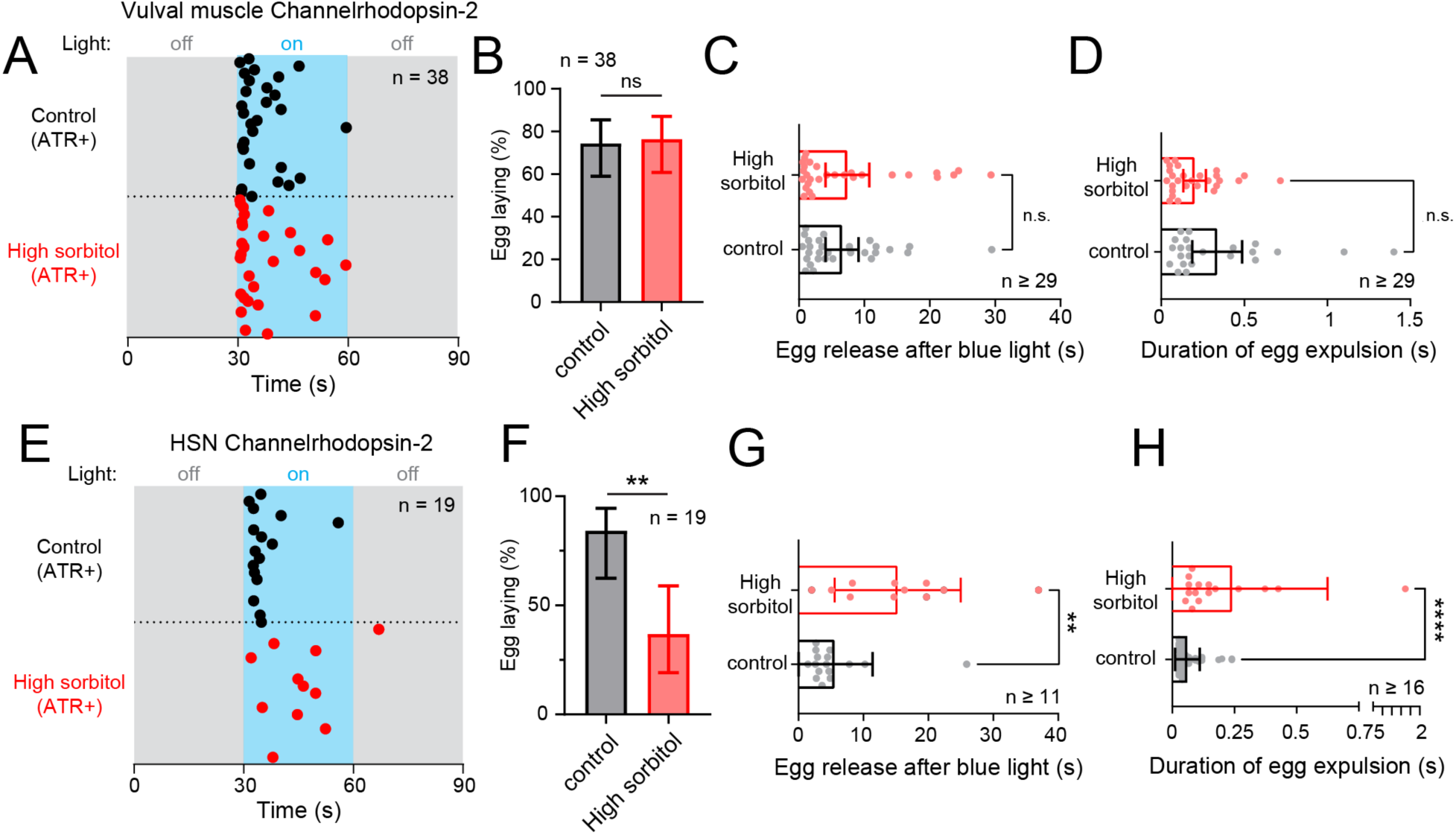
Optogenetic induction of egg laying causes delayed egg-laying responses from animals exposed to high osmolarity. **(A)** Egg-laying pattern following optogenetic activation of the vulval muscles. Dots indicate timing of the first egg-laying event. Grey shaded regions represent blue light off while blue shaded region represents blue light on. **(B)** Percent of animals showing egg laying following optogenetic stimulation of the vulval muscles. n.s. indicates not significant (p=value >0.05; Fisher’s exact test). **(C)** Bar graph with scatterplot showing mean time taken for first egg-laying event since blue light activation of the muscles. n.s. indicates not significant (p=value >0.05; Mann-Whitney test). **(D)** Bar graph with scatterplot showing mean time taken for eggs to be expelled from uterus to outside environment. n.s. indicates not significant (p=value >0.05; Mann-Whitney test). **(E)** Egg-laying pattern following optogenetic activation of the HSN neurons; as in (A). **(F)** Percent of animals showing egg laying following optogenetic stimulation of the HSNs. Asterisks ** indicate p-value = 0.0069 (Fisher’s exact test). **(G)** Bar with scatterplot showing mean time taken for egg-laying events after blue light activation of the HSNs. Asterisks ** indicate p-value 0.0022 (Mann-Whitney test). **(H)** Bar graph with scatterplot showing mean time taken for egg expulsion during egg-laying event after blue light activation of the HSNs. Asterisks **** indicate p-value <0.0001 (Mann-Whitney test). Error bars represent ±95% confidence interval for the mean.

We repeated the experiment using a transgenic strain expressing Channelrhodopsin-2 in the HSNs (Emtage et al., 2012; Gürel et al., 2012). This strain shows robust light-induced egg laying on NGM control plates (**Figure 5E**), but fewer of the animals showed egg release (∼40%) in response to continuous blue-light stimulation on high sorbitol (**Figure 5F**). Surprisingly, on high sorbitol plates, there was a delay on the onset of egg release with the first egg being laid after 10 s compared to ∼4 s for animals on NGM control media alone (**Figure 5G**). We also found that each egg physically took longer to be released on hyperosmotic media (**Figure 5H**) with egg passage occurring over 250 msec on high sorbitol rather than 50 msec seen on NGM control media, suggesting that the kinetics of egg release is sensitive to the hydrostatic pressure gradient. Together, these results suggest that hyperosmotic inhibition may prevent the normal activation of the vulval muscles for egg laying and/or efficient egg release. High sorbitol reveals deficiencies in either the optogenetic activation of the HSNs themselves or the downstream responses of the vulval muscles that drive egg laying.

To examine directly how changes in the osmotic pressure gradient affects vulval muscle activity and/or egg release, we measured vulval muscle Ca^2+^ following optogenetic stimulation of the HSNs or vulval muscles (**Figure 6A**). We moved animals from NGM to either 100 mM or 400 mM sorbitol plates for at least 1 hr prior to imaging with all animals incubated on low or high osmolarity media for ≤3 h. Under these conditions, optogenetic stimulation of the HSNs in animals on either 100 mM or 400 mM sorbitol caused a robust induction of vulval muscle Ca^2+^ transients and egg laying (≥85% of animals) during a 90 s recording period (**Figure 6B**). We observed no significant differences in the frequency of induced vulval muscle Ca^2+^ transients or egg laying in animals treated with 400 mM sorbitol (**Figure 6C**), although we did observe a significant increase in the amplitude of Ca^2+^ transients during both egg-laying (**Figure 6D**) and twitch transients (**Figure 6E**). This suggests either high osmolarity increases the vulval muscle Ca^2+^ response or that increased Ca^2+^ responses are necessary to drive vulval muscle contraction following osmotic adaptation. Our previous HSN ChR2 experiments showed high osmolarity inhibited egg release–a result not observed in these Ca^2+^ imaging experiments. The source of this discrepancy is not clear, although it may relate to differences in media composition or how the HSNs are optogenetically stimulated with pulsed illumination being used for GCaMP5 imaging rather than continuous blue light exposure typically used in behavior experiments.

**Figure 6:**
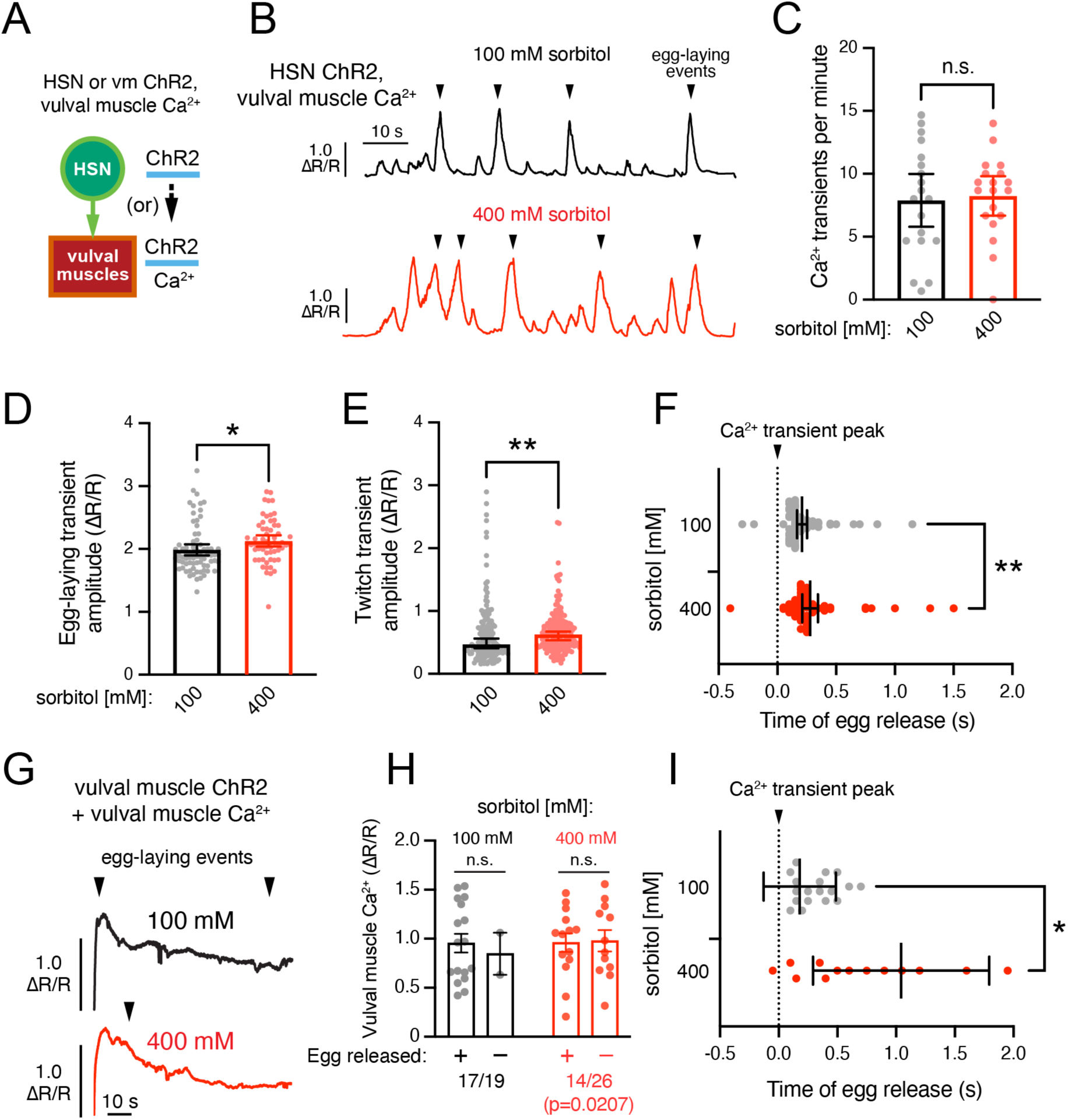
High osmolarity slows egg release following vulval muscle contraction. **(A)** Cartoon of experiment. Channelrhodopsin-2 (ChR2) is transgenically expressed either in the HSNs or vulval muscles for optogenetic stimulation and GCaMP5 and mCherry are expressed in the vulval muscles for ratiometric Ca^2+^ imaging following ∼1 h shift to agar plates containing 100 mM or 400 mM sorbitol. **(B)** Representative vulval muscle Ca^2+^ traces following optogenetic HSN stimulation. Egg-laying events are indicated by arrowheads. **(C)** Average frequency of vulval muscle Ca^2+^ transients following optogenetic stimulation. Dots indicate the per recording average with n=19 animals measured per condition. Error bars indicate ±95% confidence intervals for the mean, and n.s. indicates p-value >0.05 (Student’s t test). **(D-E)** Average vulval muscle Ca^2+^ transient amplitudes following HSN optogenetic activation during egg-laying (D) and twitch (e.g. non-egg-laying, E) events. Dots indicate single transient amplitudes pooled from 19 animals per condition. * in D indicates p-value=0.0285 (Student’s t test); ** in E indicates p-value =0.0034 (Mann-Whitney test). (**F**) Lag time between vulval muscle Ca^2+^ transient peak (dotted line) and egg release. Dots indicate timing of individual egg-laying events pooled from 19 animals. Error bars indicate 95% confidence intervals for the mean; ** indicates p-value =0.0067 (Mann-Whitney test). **(G)** Representative vulval muscle GCaMP5/mCherry fluorescence traces of Ca^2+^ activity following optogenetic vulval muscle stimulation. Egg-laying events are indicated by arrowheads. **(H)** Vulval muscle Ca^2+^ transient amplitudes following vulval muscle optogenetic stimulation. Dots indicate Ca^2+^ transient amplitudes immediately prior to egg release (plus). For those traces where no egg-laying was observed (minuses), dots indicate the peak Ca^2+^ transient amplitude from each recording. Error bars indicate standard error for the mean; n.s. indicates p-value >0.05 (one-way ANOVA with Bonferroni correction for multiple comparisons). Fraction of animals showing egg-laying events following optogenetic stimulation was significantly different (Fisher exact test, p-value =0.0207; 100 mM sorbitol: n=19 animals, 400 mM sorbitol: n=26 animals). **(I)** Lag time between vulval muscle Ca^2+^ transient peak (dotted line) and egg release. Dots indicate timing of individual egg-laying events relative to each Ca^2+^ transient peak. Error bars indicate 95% confidence intervals for the mean; * indicates p-value =0.0156 (Mann-Whitney test).

We have previously shown that eggs are released immediately after the vulval muscles reach their peak Ca^2+^ activity (Collins et al., 2016; Medrano and Collins, 2023). To measure if high osmolarity affects the coordination of vulval muscle Ca^2+^ activity and egg release, we measured how long it took for each egg to be released after the vulval muscle Ca^2+^ transient peak was observed. As shown in **Figure 6F**, in animals grown on 100 mM sorbitol, this lag between Ca^2+^ peak and egg release was ∼0.2 s on average (median 0.15 s). Animals on 400 mM sorbitol showed a significant delay to ∼0.28 s (median 0.2 s) between vulval muscle Ca^2+^ peak to egg laying. While this increased lag on 400 mM sorbitol represents a delay of only 1 frame (0.05 s), it is statistically significant (p-value =0.0067, Mann-Whitney test). Together, these results suggest changes in osmolarity affect the hydrostatic pressure gradient and the timing of egg ejection following vulval muscle contraction.

We separately compared how changes in osmolarity affected vulval muscle Ca^2+^ activity and egg laying following direct optogenetic stimulation. As shown previously (Kopchock et al., 2021), optogenetic stimulation of the vulval muscles drives an immediate and robust vulval muscle Ca^2+^ activity and egg laying (**Figure 6G**). We found no significant difference in the pattern or peak amplitude of vulval muscle Ca^2+^ responses in animals on 100 mM or 400 mM sorbitol following (**Figure 6H**). This was surprisingly because we found that animals on 400 mM sorbitol laid eggs only half as frequently as those on 100 mM sorbitol (**Figure 6H**), similar to that seen previously following optogenetic stimulation of the HSNs (**Figure 5F**). This led to a significant increase in the lag observed between the vulval muscle Ca^2+^ peak and egg release (**Figure 6I**). In animals grown on 100 mM sorbitol, the lag between Ca^2+^ peak and egg release was ∼0.18 s on average (median 0.25 s), similar to that seen following HSN optogenetic stimulation (compare **Figures 6F to 6I**). However, animals on 400 mM sorbitol showed a significant delay to ∼1 s (median 0.675 s) between vulval muscle Ca^2+^ peak to egg laying following optogenetic vulval muscle stimulation (**Figure 6I**). These results again suggest vulval muscle contraction is insufficient for egg release in animals where the hydrostatic pressure has been disrupted by high osmolarity. Egg laying therefore has two components: first, vulval muscle Ca^2+^ activity drives contraction and vulval opening; second, a high-inside hydrostatic pressure gradient powers egg release into the environment.

## Discussion

In this study we have investigated roles for both sensory and biophysical pathways in the inhibition of *C. elegans* egg-laying behavior by high osmolarity. Chemosensory mutants with defects in the detection of high osmolarity continue to lay eggs in high sorbitol, suggesting the well-understood sensory pathway mediating osmo-avoidance is not mediating the egg-laying inhibition. However, mutants that produce excessive glycerol – which would presumptively maintain the animals’ hydrostatic pressure gradient – show increased egg laying, resuming egg laying more quickly upon acute shifts to hyperosmolarity. Egg laying in animals cultured under chronic hyperosmotic conditions also show resistance. Intriguingly, such animals show an acute increase in egg laying when shifted back to hypo-osmotic conditions, consistent with the idea that the hydrostatic pressure gradient, not long-term adaptive changes, underlie the egg-laying responses. Resumption of egg laying after chronic exposure to high osmolarity does not quite restore the normal rate of vulval muscle Ca^2+^ activity nor egg laying, possibly because of additional long-term effects on egg production overall (**Figure 1G, right**). However, optogenetic stimulation of either the HSNs or the vulval muscles is sufficient to induce vulval muscle Ca^2+^ activity and egg release even when egg laying is otherwise inhibited by high osmolarity conditions. Interestingly, high osmolarity slows egg release following vulval muscle Ca^2+^ activity. Taken together, these results suggest that both a sensory pathway and a biophysical pathway act in parallel to mediate inhibition of *C. elegans* egg-laying behavior in hyperosmotic environments.

Support for the idea that high osmolarity can inhibit egg laying via two distinct pathways is bolstered by recent results showing that increasing concentration of sorbitol inhibit egg laying (Huang et al., 2023). They find that animals with mutations in *tax-2* or *tax-4* can lay eggs in 300 mOsm conditions, while wild type animals are unable to. Interestingly, these same mutants are unable to lay eggs at higher concentrations (375 or 450 mOsm), as we observe (at 400 mOsm), suggesting a separate pathway for inhibition via higher osmotic conditions. Our data for a biophysical mechanism is consistent with this secondary inhibitory pathway. Separate pathways in the detection of a physical stimulus have been observed in the *C. elegans* gentle and harsh touch response where different mechanosensors and mechanosensory neurons are responsible for detecting and conveying information on the physical stimulus depending on the intensity of that stimulus (Chalfie, 2014; Fechner and Goodman, 2018; Goodman, 2006). Likewise, it is possible that, at the high osmotic conditions tested in this study (400 mM sorbitol), a mechanism that inhibits egg-laying independent of chemosensation exists. We propose a parallel biophysical model in which acute sensory detection of high osmolarity signals to inhibit egg laying, but, under conditions of high or chronic osmotic conditions, lack of osmosensory detection cannot bypass or overcome the biophysical control of egg release. In this case, egg laying is physically inhibited by a nonpermissive osmotic pressure barrier that regulates a hydrostatic pressure gradient across the cuticle that supports egg release upon vulval opening. Such a model would predict a physical limit at which the osmotic pressure becomes too great for egg laying to occur. In support of this, we observe that animals acclimated to or maintained on 400 mM sorbitol over multiple generations show a steady, but lower, level of egg laying when compared to animals maintained in 100 mM sorbitol media (**Figures 1 and 3**). This suggests that egg release occurs only when a sufficient buildup of pressure, perhaps through increased egg accumulation (**Figure 1G**), is present to overcome either physical limitations and/or the inhibition of circuit activity that accompanies osmosensory input. Future studies looking at whether acute increases in internal pressure, such as microinjection (Medrano and Collins, 2023), can bypass osmotic inhibition of egg laying may provide further evidence to the biophysical blocking of egg laying model.

A prediction of our biophysical model is that hyperosmotic inhibition of egg laying acts independent of egg-laying circuit activity. In other words, animals on high osmolarity would show normal levels of Ca^2+^ activity in the egg laying circuit but would simply be unable to physically lay eggs. Although we observe a steady (albeit reduced) rate of egg laying from animals acclimated and maintained in high osmolarity (**Figure 3C and D**), vulval muscle Ca^2+^ activity is reduced in animals exposed to 400 mM sorbitol for up to 4.5 hours. (**Figure 4D and E**). This supports past work that suggests that sensory signaling of high osmolarity environments signals to inhibit egg-laying activity (Huang et al., 2007; Zhang et al., 2008). However, what happens to the circuit activity of the egg-laying circuit (the HSNs, VCs, uv1s, and vulval muscles) after long-term growth on high osmolarity media remains unclear. We hypothesize a return of the same pattern of two-state circuit activity that accompanies the onset of the egg-laying active state in normal osmotic conditions (Collins et al., 2016; Ravi et al., 2018b; Waggoner et al., 1998). Thus, future studies should focus on investigating the long-term changes in the patterns of Ca^2+^ activity of animals chronically exposed to hyperosmotic environments. The use of sensory mutants should allow direct tests of the distinct contributions and mechanisms of the two pathways of hyperosmotic inhibition (Zhang et al., 2022).

Can animals that are adapted to high osmolarity via consistent production of glycerol bypass osmolarity-mediated inhibition of egg laying? *osm-7* and *osm-8* mutants are still inhibited for egg laying at 400 mM sorbitol (**Figure 2C**). As such, sensory signaling may still be acting to depress circuit activity and prevent egg laying. However, our experiments do not account for possible increased rates of glycerol excretion in *osm-7/8* mutants. Upon movement from hyperosmotic to isotonic environments (now relatively hypoosmotic to hyperosmotic-acclimated animals), *C. elegans* shows a rapid efflux of glycerol, losing about 25% of accumulated glycerol within a few hours (Lamitina et al., 2004). This efflux is likely triggered via swelling and mediated by the glycerol-permeable aquaglyceroporins, AQP-1 and AQP-7 (Huang et al., 2007; Lamitina et al., 2004). As such, it is possible that *osm-7* and *osm-8* mutants have increased baseline levels of glycerol excretion to acclimate to relative hypotonic environments, which would typically be isotonic to animals lacking such mutations. In this scenario, *osm-7/8* mutants would thus require further acclimatization once they are shifted to higher osmolarity conditions, possibly explaining why we still see inhibition of egg laying in high osmolarity from these mutants. However, we do see that *osm-8* mutants resume egg laying much more quickly after shifting to 400 mM, consistent with glycerol production in these animals conferring resistance, at least in part. Future studies investigating if double mutants of *osm-7/8* and aquaglyceroporins can lay eggs in high osmolarity will help reveal whether overcoming a hydrostatic gradient is sufficient to lay eggs in high osmolarity.

A rapid excretion of glycerol also helps explain our observations that moving animals from high osmolarity to a relatively lower osmolarity induces an increase in egg laying that occurs only within the first 30 minutes. This is because, similarly, shifting animals from high to lower osmolarity causes an acute initial increase in relative body size, followed by a return to a typical size, that occurs within 20 minutes and is likely mediated by glycerol or water excretion (Huang et al., 2007; Lamitina et al., 2004). Consistent with a model in which an acute increase in internal pressure induces egg-laying, the sudden influx of water would then lead to an increase in hydrostatic pressure and egg-laying behavior, as we observe (**Figure 3**). Past work has shown that acute shifts to ‘low’ external osmolarity promotes egg-laying circuit activity (Zhang et al., 2008), and our results would predict that any shift from a relatively higher to lower osmolarity would increase the steepness of the ‘high inside’ hydrostatic pressure gradient, inducing egg-laying circuit activity and behavior. Indeed, this interpretation is consistent with our results showing artificial egg accumulation or even acute injection drives a robust activation of egg-laying circuit activity and behavior (Medrano and Collins, 2023). Thus, future studies should focus on investigating whether induction of egg-laying circuit activity is due to a signaling pathway or to the influx of water causing an acute stretch (or both). Measurements of circuit activity during these acute shifts would also help reveal the mechanisms by which egg laying is activated or inhibited.

We find that optogenetic stimulation of the vulval muscles drives robust egg laying behavior even in animals on high osmolarity, but stimulation of the HSNs under these conditions leads to defects in egg laying. As optogenetic activation of the HSNs is normally sufficient in driving egg-laying active states in norm-osmic conditions (Collins et al., 2016), our results suggest that hyperosmotic inhibition increases the activity threshold required to drive egg-laying behavior. These are consistent with results showing that optogenetic HSN stimulation was only able to show a robust induction of vulval muscle Ca^2+^ activity when there were sufficient eggs in the uterus, with egg accumulation elevating the hydrostatic pressure gradient (Ravi et al., 2018b). Our results further suggest that inhibition can be bypassed by direct activation of the vulval muscles, as we see using acute injection which activates the vulval muscles (Medrano and Collins, 2023). Recent evidence suggest multiple subthreshold signals are required to drive proper vulval muscle activity and egg-laying behavior of *C. elegans* (Olson et al., 2023). Given that high osmolarity inhibits both VC and HSN activity (Zhang et al., 2008), our results in which the HSNs fail to drive robust egg laying are consistent with this multiple input model. This further suggests a general mechanism by which inhibitory stimuli converge to depress activity in several cells of a circuit including the HSNs (Ravi et al., 2021), reducing the probability that Ca^2+^ activity in the circuit will drive a given behavior. How are the HSNs inhibited by high osmolarity? Past work has suggested that TRPV channels expressed on the uv1s may signal to inhibit egg-laying activity in high osmolarity as *ocr-2(vs29)* mutants show normal levels of egg laying (Zhang et al., 2010). However, we fail to replicate these results precisely and find *ocr-2(vs29)* mutants are still inhibited for egg laying in high osmolarity (**Figure 2D**). A possible confound of these results is the strongly reduced brood size of *ocr-2(vs29)* mutants which could complicate accurate measurements of egg-laying inhibition. Alternatively, *flp-17* neuropeptide signaling from BAG neurons has been recently proposed to mediate inhibition of egg laying via detection of hyperosmotic environments. BAG ablated and *flp-17* mutants show normal egg laying on 300 mOsm conditions (Huang et al., 2023), but whether this is also true for inhibition by higher osmolarity remains unknown. As such, more studies identifying conditions where egg laying occurs even in hyperosmotic environments would help reveal the contributions of the chemosensory and biophysical pathways for osmolarity-driven inhibition of egg laying.

## Acknowledgements

We thank Dr. James Baker and members of the Collins Lab for constructive feedback on the manuscript.

## Competing interests

The authors report no competing interests.

## Funding

This work was funded by grants from the National Institutes of Health (NS086932) and the National Science Foundation (IOS-1844657) to K.M.C.; E.M. was supported by a National Science Foundation Graduate Research Fellowship and McKnight Doctoral Fellowship. Some strains were provided by the *C. elegans* Genetics Center, which is funded by NIH Office of Research Infrastructure Programs (P40-OD010440).

## Data availability

Raw data will be made available through FigShare.

## Notes

### Competing Interest Statement

The authors have declared no competing interest.

